# Effects of multistability, absorbing boundaries and growth on Turing pattern formation

**DOI:** 10.1101/2024.09.09.611947

**Authors:** Martina Oliver Huidobro, Robert G. Endres

## Abstract

Turing patterns are a fundamental concept in developmental biology, describing how homogeneous tissues develop into self-organized spatial patterns. However, the classical Turing mechanism, which relies on linear stability analysis, often fails to capture the complexities of real biological systems, such as multistability, non-linearities, growth, and boundary conditions. Here, we explore the impact of these factors on Turing pattern formation, contrasting linear stability analysis with numerical simulations based on a simple reaction-diffusion model, motivated by synthetic gene-regulatory pathways. We demonstrate how non-linearities introduce multistability, leading to unexpected pattern outcomes not predicted by the traditional Turing theory. The study also examines how growth and realistic boundary conditions influence pattern robustness, revealing that different growth regimes and boundary conditions can disrupt or stabilize pattern formation. Our findings are critical for understanding pattern formation in both natural and synthetic biological systems, providing insights into engineering robust patterns for applications in synthetic biology.

**Author summary:** During development, tissues self-organize to go from a single cell to a structured organism. In this process, simple chemical reactions lead to the emergence of the intricate designs we see in nature, like the stripes on a zebra or the labyrinths on a brain cortex. Although multiple theories have been proposed to model this phenomenon, one of the most simple and popular ones was introduced in the 1950s by the mathematician Alan Turing. However, his theory oversimplifies the biological conditions and ignores properties such as non-linearities, boundary effects, or growth in the tissue. In this work, we used a combination of mathematical models and computer simulations to investigate how these real-world factors influence pattern formation. Our findings show that when we account for these realistic effects, the patterns that emerge can be very different from what Turing’s theory would predict. Thus, this work may help us better understand the laws behind pattern formation and could have practical applications in tissue engineering for medical or environmental applications.

## Introduction

How biology produces robust patterns in space and time is still largely an open question [1]. In 1952, Alan Turing proposed a mechanism referred to as Turing patterns or diffusion-driven instabilities, which explains how a homogeneous tissue results in self-organized spatial repetitive patterns of the network’s molecules [2, 3]. However, this proposed mechanism is far from real biological complexity, and suffers from fine-tuning, meaning only a small subset of constrained parameters can generate these spatial patterns [4, 5]. Additionally, the diffusion-driven instability is based on linear stability analysis which often does not address realistic phenomena such as multistability, growth, exotic boundary conditions, and non-linearities [6]. A way forward to understanding the relationship between theoretical Turing patterns and real biological patterns is to engineer these reaction-diffusion networks in bacterial colonies using synthetic biology [7, 8]. This would allow us to better understand the role of Turing patterns in biological pattern formation, as well as to engineer synthetic patterns for industrial applications [9–11]. This approach was recently extended and implemented in the literature [12], where a Turing gene circuit was introduced in growing *E*.*coli* colonies producing fluorescent periodic spatial patterns. Fig 1A,B shows a final snapshot of a bacterial colony with periodic patterns in the GFP channel and the mean fluorescence in a cross-section of the colony. Understanding how patterns arise in this biological system is crucial for engineering synthetic patterning and to further understand the mechanisms of pattern formation in biology.

**Fig 1.**
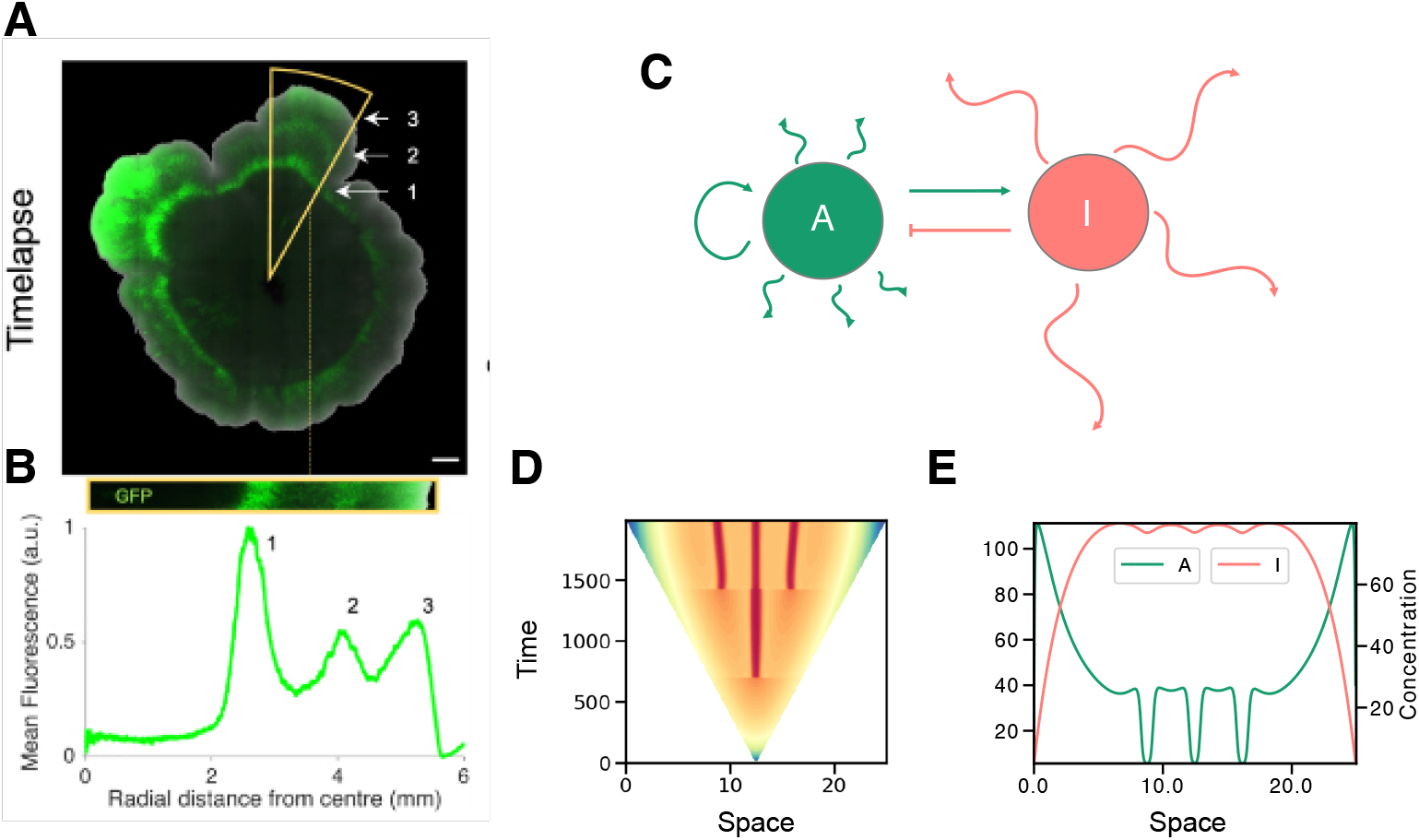
Synthetic Turing patterns from experiments and theory. A,B subfigures are generated by J.Tica and D.Bazzoli and published in [12] **(A)** Last frame of a time-lapse of a bacterial colony grown with a synthetic 6-node Turing gene circuit producing periodic spatial patterns in two-dimensions (scale bar: 1mm). Only the GFP channel is shown. **(B)** Mean fluorescence radial profile of the colony along the highlighted wedge with azimuthal averaging (yellow triangle). **(C)** Two-node network based on Turing’s original paper [2] with slow diffusing activator *A* in green and fast diffusor inhibitor *I* in pink. **(D)** Kymograph of numerical solution for *A* in 1D of the Turing circuit shown in C (corresponding to activator *A*). The domain is growing with absorbing boundaries. Plotted as a time-series of molecule *A* (nM). The *Y* axis is time (h), the *X* axis is space (mm) and concentration is shown by color with blue being high concentration and red being low concentration. A periodic pattern appears as the tissue grows. **(E)** Final snapshot of the solution in D shows a periodic pattern. The *Y* axis is the concentration (nM) of *A* (left, green) and *I* (right, pink) while the *X* axis is space (mm).

A realistic mathematical description of such a biological system has non-linearities, multistability, growth, and boundary conditions which are not adequately captured with linear stability analysis. The latter technique only describes the onset of pattern formation, not the final pattern. Several studies have explored how these phenomena affect Turing pattern formation including non-linearities [13], multistability [6], boundaries [14–19] and growth [20–22]. However, these studies are often based on idealized domains, certain types of popular or convenient boundary conditions, and artificial growth assumptions. Additionally, often they do not provide statistics of how Turing patterns become more or less robust to the fine-tuning problem when introducing these phenomena by conducting high-throughput parameter scans. Therefore, a high-throughout numerical study is needed to understand how all these natural phenomena affect Turing pattern formation, in particular in a synthetic system such as the one in [12].

In this paper, we combine linear stability analysis and numerical solutions, to study a simple Turing reaction-diffusion network (Fig 1C) with non-linearities which leads to multistability. Our analysis goes beyond classical Turing patterns as we also consider other types of instabilities such as Hopf and Turing I Hopf and how these can generate periodic stationary or non-stationary patterns. We find that switching of steady states in multistable systems can generate unexpected outcomes not predicted by linear stability analysis. Additionally, growth and realistic boundary conditions are added to understand how patterning occurs in microbial colonies with synthetic Turing gene circuits. This type of model results in patterns in growing domains such as those seen in Fig 1D,E which resemble the biological example in Fig 1A,B. We find that different boundaries and growth regimes can break or form patterns, therefore adding or removing robustness to pattern formation. Studying how all these biological phenomena affect patterning is extremely important not only in the context of engineering patterns in synthetic biology, but also to understand how robust patterns occur in developmental biology where large gene-regulatory networks commonly exhibit non-linearities and multistability, and where tissues grow while in contact with an external environments.

## Results

To study the effects of multistability, non-linearities, boundary conditions, and growth, we use a simple reaction-diffusion model which consists of a 2-node Turing topology system with non-linear Hill functions to represent gene activation and inhibition with cooperativity. The topology can be seen in Fig 1C where the activator *A* activates itself and the inhibitor *I*, while the inhibitor *I* inhibits the activator *A*:

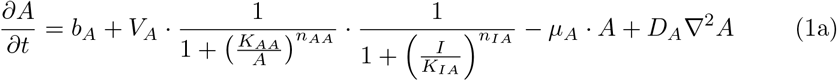

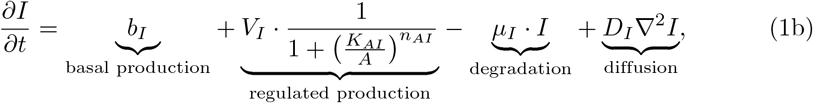

with *X* being the concentration of species *X* = *A, I* and the Laplacian ∇^2^ = *∂*^2^*/∂x*^2^ in one spatial dimension (1D). The differential equations describe the rate of change in space and time as a combination of basal production and regulated production, as well as degradation and diffusion. Parameter *b*_*X*_ is the basal production rate which corresponds to the leakage of the promoter, producing molecules even when the system is fully inactivated. Furthermore, *V*_*X*_ is the induced maximum production rate, *K*_*XY*_ is the dissociation constant of *X* binding to *Y* ‘s regulatory DNA, *n*_*XY*_ is the Hill coefficient (cooperativity constant) of *X* binding to *Y* ‘s regulatory DNA, and *µ*_*X*_ is the linear degradation rate of *X*. Finally, *D*_*X*_ is the diffusion constant of species *X*.

The boundary conditions and domain are not strictly defined here as they vary throughout the study. The boundary conditions used are Neumann for reflective boundaries, where the derivative at the boundary is zero, as well as Dirichlet for absorbing boundaries, where the value at the boundary is zero. The domain used is either a fixed domain of length *L* or a linearly, isotropically and apically growing domain, which replicates the experimental growth in 1D of the bacterial colonies in [12] (or Fig 1A).

### From analytical to numerical: other types of dispersion relations and patterns

Turing instabilities are defined using linear stability analysis (LSA) as systems that are stable without diffusion but unstable for a finite wave number as diffusion is introduced (see Methods). These Turing instabilities generally lead to stationary periodic patterns (see Fig 2B). In this section, we demonstrate how sometimes the classical Turing instability theory fails to predict pattern formation. Other types of dispersion relations beyond classical Turing instabilities can produce stationary patterns and non-stationary regular patterns that might be of interest in developmental biology.

**Fig 2.**
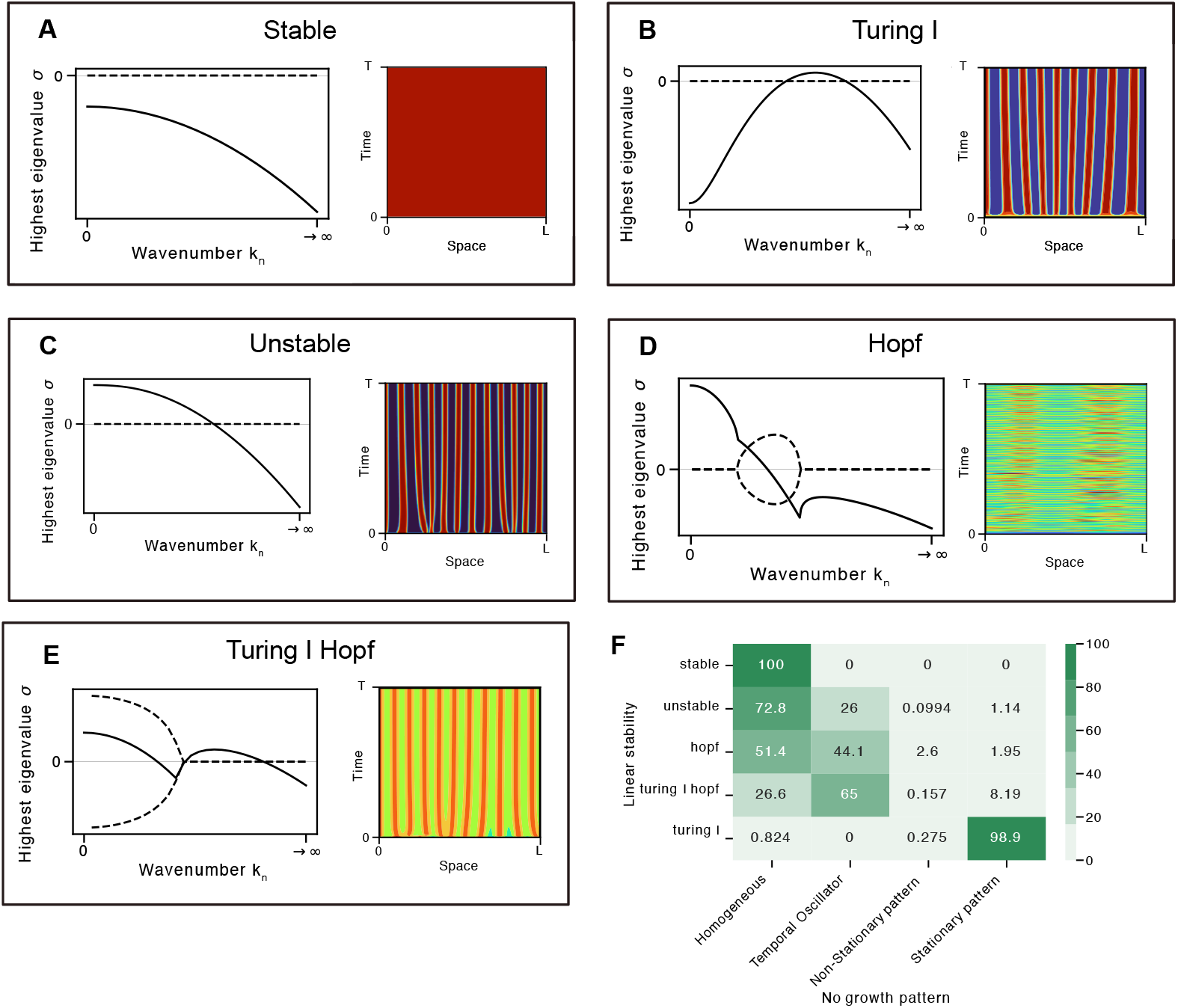
Relationship between dispersion relation and numerical solution in mono-stable systems. **(A-E)** Examples of dispersion relations (left) and resulting numerical solutions (right) for Stable, Turing I, Unstable, Hopf, and Turing I Hopf. **(F)** Confusion matrix linking outcomes from LSA (rows) and numerical calculation (columns). Numbers show the percentage of solutions across the LSA output rows.

High-throughput studies such as [5, 23, 24] only consider the Turing-I type dispersion for patterning and disregard the rest. Here, we study solutions of the classical 2-node circuit (Fig 1C, Eq 1) and explore beyond Turing I and stationary patterns to give a more complete view of the relationship between LSA and spatio-temporal patterns. The kinetic and numerical parameters used are shown in Tables Table S1 and Table S2.

First, LSA was carried out for particular parameter sets to find multiple steady states with different types of stability. Second, numerical simulations were computed using the Crank–Nicolson algorithm, where the initial condition is drawn from a random uniform distribution around a particular steady state. By classifying dispersion-relation and pattern types, we document what type of dispersion relations in mono-stable systems can be linked to what types of patterns, to gain insights into predicting pattern formation from LSA. All the solutions used in this section are mono-stable systems to ensure a direct correlation between the dispersion relation and the pattern obtained. Details on LSA, Crank-Nicolson and pattern classification can be found in the Methods section.

All types of dispersion relations in our parameter space were classified into 5 types: stable dispersion relations have all eigenvalues *σ* below zero for any wavenumber *k*_*n*_ (Fig 2A). Unstable dispersion relations have a positive eigenvalue at *k*_0_ = 0 which eventually drops below zero as diffusion is introduced, i.e. *k*_*n*_ *>* 0 (Fig 2C). Hopf-type dispersion relations, as with any unstable dispersion relation, show an instability without diffusion (*σ >* 0 for *k*_0_ = 0), which eventually drops below zero for positive wavenumbers. However, in the case of the Hopf-type dispersion relation, when the eigenvalues cross the zero line, there is a pair of complex conjugate eigenvalues (Fig 2D). A Hopf-type dispersion relation is different from a Hopf bifurcation: a bifurcation displays a shift in stability as a model parameter changes, while the Hopf-type dispersion is a change in stability as a function of the wavenumber *k*_*n*_. Turing I dispersion relations, as previously mentioned, are stable without diffusion, have an instability for a positive wavenumber, and finally become stable again for very large wavenumbers (Fig 2B). Turing I Hopf dispersion relations, are a combination of Turing I and Hopf. As the Hopf-type dispersion, they are unstable without diffusion. Then, as *k*_*n*_ is increased, the system becomes stable with a pair of complex conjugates as the eigenvalues cross the zero line. Finally, a Turing I-type behavior arises getting a peak above zero and decaying again for large wavenumbers (Fig 2E).

Other types of dispersion relations exist, which are not displayed here such as Turing II. Unlike Turing I, the eigenvalues do not become stable again for very large wavenumbers. Therefore, this system displays an instability at very large wavenumbers, which results in infinitesimally small wavelength patterns (at least for an infinite system). These are considered to produce spatially homogeneous solutions, except in the case of space discretization where they can produce small wavelength patterns [25]. However, Turing II solutions are not possible in systems such as this one where all nodes are diffusing. This is because for *k*_*n*_ → ∞, all eigenvalues *σ* must be negative (see Eq 6).

As with the classification of the dispersion relations, we classify the patterns produced numerically into homogeneous, temporal oscillator, non-stationary, and stationary patterns (see Methods and Fig S2). By classifying both LSA and numerical outputs, we can generate a confusion matrix linking the different types of dispersion relations with the different types of spatio-temporal patterns (Fig 2F). This confusion matrix shows that also other systems than Turing I can generate stationary spatial patterns. Unstable, Hopf and Turing I Hopf can produce such results too as seen in Fig 2C-E. Additionally, interesting behaviours can arise such as temporal oscillators and non-stationary patterns (Fig 2D).

### Feedbacks and non-linearities cause multi-stability

Multi-stable systems are another case where linear stability analysis (LSA) fails to correctly predict pattern formation. In this section, we demonstrate how LSA is neither sufficient nor necessary to predict Turing patterns in multi-stable systems. In particular, we study in detail their dynamical behavior, which can lead to the creation or breaking of patterns. The motivation behind this arises from the high degree of multi-stability exhibited by biological systems, especially in those systems with non-linearities and feedback loops, which often occur in biology or synthetic systems [26, 27].

Using the two-node non-linear Turing topology (Eq 1), multi-stable solutions were identified by finding the steady states of the system using the Newton-Raphson method. These multi-stable solutions were then studied to understand how the patterning dynamics are affected in the presence of multiple steady-state solutions. As in the previous section, LSA was carried out to find multi-stable solutions and the Crank–Nicolson algorithm was run to obtain the numerical solution for these. Following the classical hypothesis used in the Turing literature, we only expected equations fulfilling the Turing conditions to produce patterns. Here, we present various examples of how this hypothesis can break in the presence of multi-stability.

Fig 3 shows a case where diffusion-driven instability conditions are not required for Turing pattern formation. The unstable state, having no Turing instability, manages to form a Turing pattern as the neighbouring Turing steady state attracts it (see dispersion relations in Fig 3B). It therefore produces a stationary pattern (Fig 3C-middle), even though its dispersion relation does not predict so. The trajectory is depicted in the phase diagram (Fig 3A) which shows the steady states along with the vector field to understand the potential trajectories of the system. The phase diagram does not fully capture the dynamics as it describes the system without diffusion, while the dispersion relation and the numerical solution do consider diffusion.

**Fig 3.**
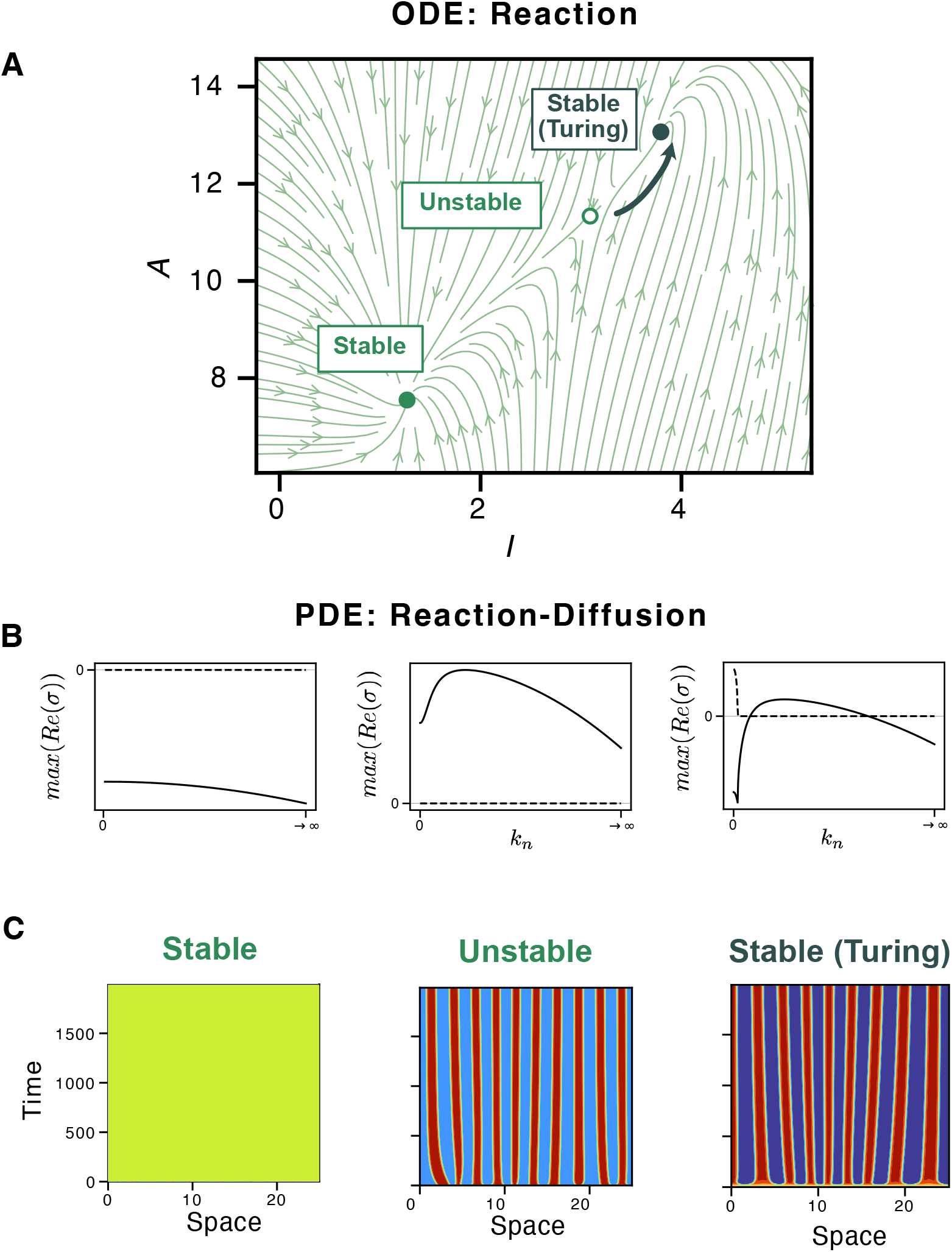
Stationary patterns with multi-stability. **(A)** Phase diagram without diffusion illustrating three distinct steady states where the derivative is zero: stable, unstable, and stable (Turing). These steady states are represented within a parameter space defined by the two concentrations for species *A* and *I*. The vector field, indicated by light green arrows, shows the direction of the derivatives of the system at various points in the parameter space. A hand-drawn trajectory is also shown (dark green arrow), demonstrating how the unstable state may evolve into the Turing state. **(B)** Dispersion relation showing each type of state. **(C)** Numerical solutions of the three steady states with diffusion, where the unstable state unexpectedly produces a Turing-like stationary pattern.

Next, we present a case where LSA incorrectly predicts stationary pattern formation. Fig 4 shows an ephemeral or transient pattern that occurs in the unstable and Turing regimes. The Turing pattern initially develops in the vicinity of the Turing steady state. As the spatial heterogeneity is amplified and settles, it is attracted by the stable steady state, leading to the disruption of the pattern. This type of transient pattern behavior was also recently reported in [6].

**Fig 4.**
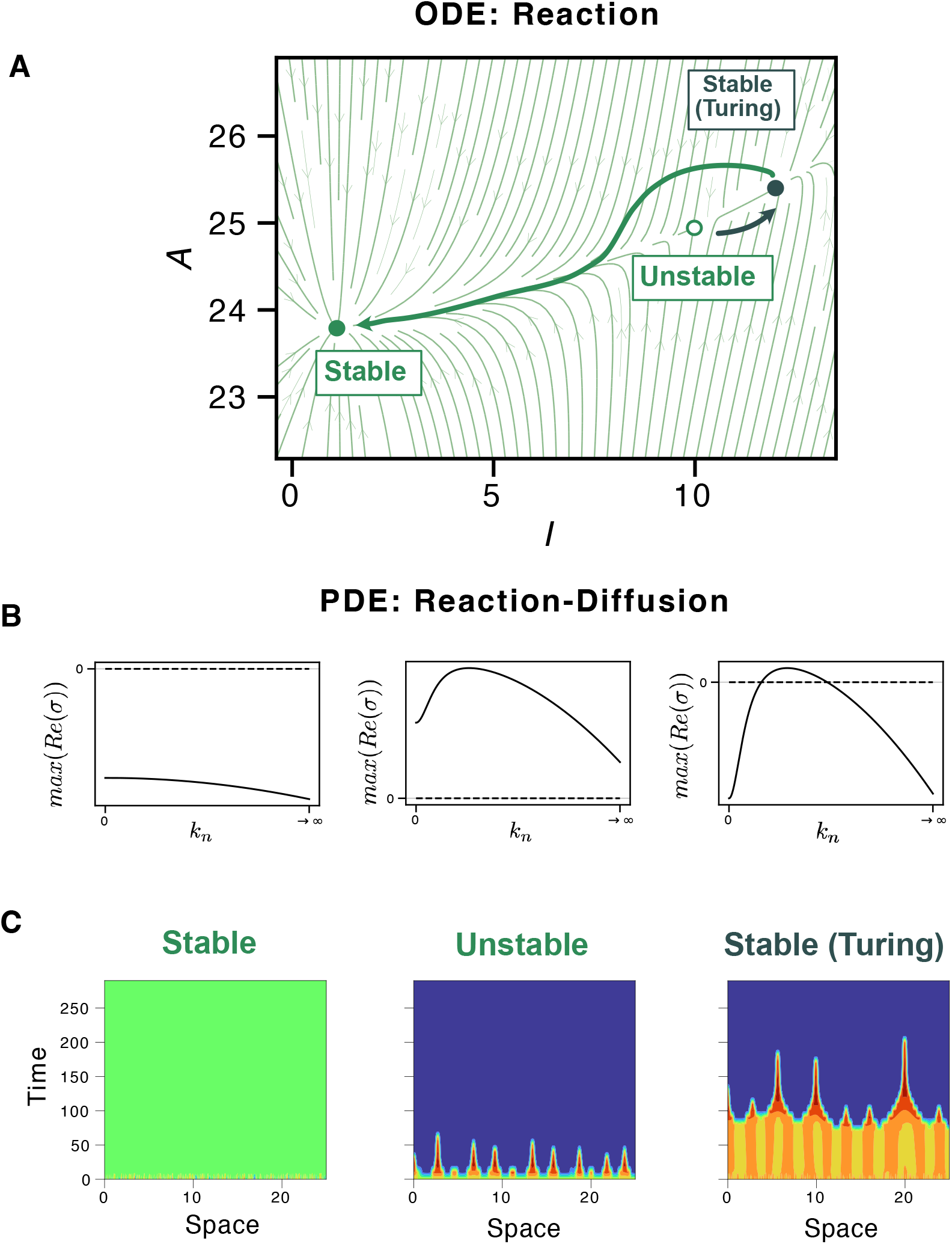
Ephemeral patterns with multi-stability. **(A)** Phase diagram without diffusion illustrating three distinct steady states where the derivative is zero. The hand-drawn trajectory (dark green arrow) shows an unstable state evolving into a stable (Turing) and a regular stable (non-Turing) state, indicated by thick black and green arrows, respectively. **(B)** Dispersion relation showing each type of state. **(C)** Numerical solutions of three steady states with diffusion. Unstable and Turing states produce temporary periodic stationary patterns that then disappear and become spatially homogeneous solutions.

Other interesting examples can be found, for example, where an unstable state is surrounded by two Turing states. This unstable state leads robustly to a Turing pattern (Fig S4A). Additionally, in some cases, the unstable state settles into Turing, but the Turing state gets pulled by the stable attractor (Fig S4B). Additionally, some systems even exhibit three solutions which are homogeneous in time and space (Fig S4C). In this case, it would be worth investigating earlier time points with more resolution, as a pattern might appear then. Similar interactions occur with multi-stability involving Turing I Hopf solutions as seen in Fig S4D.

### Biological features: absorbing boundaries and growth

As shown above, both numerical solutions and multi-stability can break the preconception that only classical Turing I systems can produce stationary periodic patterns. Here, we look deeper into how other aspects linking the theory closer to the biological reality can also break this preconception. In particular, we look at how adding absorbing boundaries and growth to a reaction-diffusion system may induce or break patterning. This particular direction was inspired by experiments described in [12], where growing bacterial colonies in agar are used as a platform to engineer Turing patterns using synthetic gene circuits (see Fig 1A,B).

The parameter space was explored using simulations, classified as (1) no-growth and reflective boundaries, (2) no-growth and absorbing boundaries, and finally (3) growth and absorbing boundaries. This way we can understand separately the effects of absorbing boundaries and growth in a synthetic system such as [12] Again, all these parameter sets have only a single steady state to ensure patterning effects are due to boundaries and growth, and not due to multi-stability. When absorbing boundaries or growth are added, periodic patterns are potentially created, disrupted, or remain the same. The Sankey diagram in Fig 5A shows how this system can transition when absorbing boundaries and growth are introduced. This transition can also be seen in Fig S5 as a confusion matrix from reflective to absorbing boundaries and from non-growing to growing domains.

**Fig 5.**
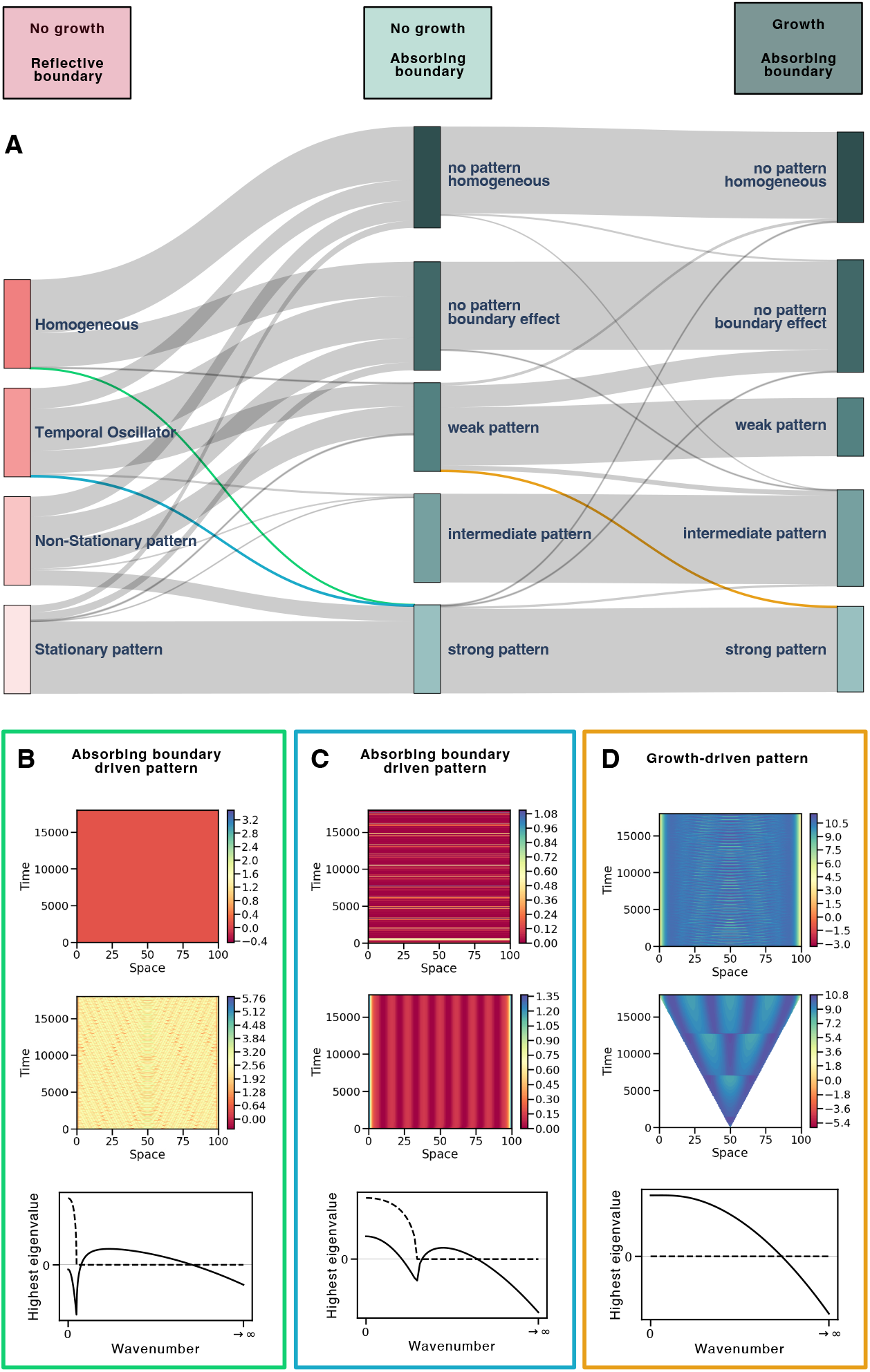
Effects of absorbing boundaries and growth on spatio-temporal patterns. **(A)** Sankey Diagram showing transitions across pattern classes when absorbing boundaries and growth are added. The lines’ thickness corresponds to the number of cases following that transition. Pink corresponds to the first classification system while green corresponds to the second classification system. Specific cases are highlighted in light green, blue, and orange. **(B)** Example of a system transitioning from a homogeneous (top) to a strong pattern (middle) upon introduction of absorbing boundary conditions. The dispersion relation (bottom) shows a Turing instability. **(C)** Example of a system transitioning from a temporal oscillator (top) to a strong pattern (middle) upon introduction of absorbing boundary conditions. The dispersion relation (bottom) shows a Turing I Hopf instability. **(D)** Example of a system transitioning from a weak pattern (top) with oscillations to a strong stationary pattern (middle) upon introduction of growing domains. The dispersion relation (bottom) shows a system

As explained in the Methods section, the classification previously used cannot be applied for absorbing boundary conditions as these prevent the pattern from being completely homogeneous or stationary. Therefore, a different type of classification has to be used for reflective and absorbing boundary conditions. For reflective boundary conditions, we classify patterns into (1) homogeneous, (2) temporal oscillators, (3) non-stationary patterns, and (4) stationary patterns (see Methods and Fig S2). For absorbing boundary conditions we classify patterns into (1) homogeneous, (2) boundary effect, (3) weak pattern, (4) intermediate pattern, and (5) strong pattern (see Methods and Fig S3). Although the classification outputs cannot be directly compared, this Sankey diagram is useful for identifying interesting cases: If absorbing boundary conditions had no effect, we would expect homogeneous and temporal oscillator categories to become no pattern, homogeneous, or boundary effect. Additionally, we would expect stationary patterns to become strong patterns. This is commonly the case as seen by the thicker lines in Fig 5A, which shows boundaries do not often affect the pattern. However, some exceptions are highlighted with color in this diagram and further studied in Fig 5B,C,D.

#### without a diffusion-driven instability

In particular, we observe systems where an absorbing boundary condition makes a homogeneous solution become a non-stationary pattern. Additionally, absorbing boundary conditions can turn a temporal oscillator into a stationary pattern. Finally, by adding growth, a non-stationary pattern can be stabilized into a growing stationary pattern.

## Discussion

Over the last few decades, there have been tremendous efforts in understanding how biology produces robust reproducible patterns, e.g. during embryonic development, with seminal work by Turing and others [2–4]. In this context, robustness refers to the fraction of parameter space leading to Turing patterns. Due to the Turing conditions, this robustness is generally tiny, hindering progress in understanding developmental patterns. Here, we investigate how a highly non-linear reaction-diffusion model motivated by synthetic gene circuits can generate spatio-temporal patterns beyond predictions from linear stability analysis (LSA). The discrepancies between LSA and numerical predictions are considerable and can be attributed to non-linearities, boundary conditions, and growth. A specific focus of our work is the role of multi-stability in Turing systems, which arises from non-linearities and feedback loops. We describe the mechanism by which an unstable system can acquire Turing patterns, and conversely how Turing states can lose their patterns. These findings make the Turing mechanism significantly more versatile than originally assumed and point towards investigating the effects of other biological properties such as absorbing boundary conditions or growing domains on patterning robustness.

Most current robustness studies primarily focus on Turing I instabilities [5, 23, 24]. However, we demonstrate that numerical methods for robustness searches can uncover a wider variety of relevant spatio-temporal solutions. For example, systems exhibiting unstable, Hopf or Turing I Hopf dispersion relations can also produce Turing-like stationary periodic patterns (see Fig 2). Therefore, systems with such dispersion relations should not be disregarded when studying pattern formation. Although considering these systems might slightly improve robustness, it does not fully explain the difference between robust mechanisms found in nature and the non-robust Turing patterns. The latter still requires fine-tuning due to mathematical constraints.

Additionally, some systems with a Hopf-dispersion relation produce noteworthy non-stationary patterns. These non-stationary patterns could be highly valuable in developmental and synthetic biology because the arrest of gene expression could transform them into stationary patterns. Furthermore, these non-stationary patterns might act as a pre-pattern for initial symmetry breaking, and make downstream patterning more reproducible. Therefore, in the context of symmetry-breaking events and patterns in developmental biology, it is crucial to consider mechanisms beyond traditional Turing dispersion relations and include Hopf, Turing I Hopf, and simple unstable systems as potential explanations for biological patterning.

In terms of multi-stability, there is evidence for the above-mentioned ephemeral patterns, i.e. transient patterns occurring as the system transitions from Turing to stable states, to be relevant for developmental biology. If the patterned state has a longer life time than down stream gene expression, then they suffice in temporarily activating the necessary genes to produce a permanent phenotype. For instance, digit formation or hair follicle development requires periodic patterns only at specific times when a hormone is produced to generate fingers or hairs [28, 29]. Understanding how interactions between different steady states influence pattern formation is essential, as multi-stability is widespread and plays a critical role in biological systems [30]. In the authors’ view, dynamics is more important than steady-state patterns for embryonic development [31].

An important finding of our work is that absorbing boundary conditions might be part of the solution to the lack of Turing patterning robustness. A significant proportion of cases fall into the top diagonal of the confusion matrix (Fig S5A), suggesting that absorbing boundary conditions might enhance patterning robustness. Specifically, absorbing boundaries can induce spatial patterning by creating non-stationary or stationary periodic patterns from spatially homogeneous patterns (Fig 5B,C).

Absorbing boundaries (and growth) were also used in the experimental setup in [12], which produced a high number of patterns despite the fragility of the Turing mechanism. However, adding absorbing boundaries can also disrupt spatial patterns, as shown in the Sankey diagram (Fig 5A). Hence, each effect is a two-sided sword in both positively and negatively affecting pattern formation.

In contrast to absorbing boundaries, introducing growth did not seem to improve robustness for pattern formation as more cases ended up at the bottom of the diagonal in the confusion matrix (Fig S5B). This finding contradicts some literature on growth-induced Turing patterns [20]. The discrepancy may arise from differences in growth rates and types of growth (e.g. exponential or logistic). Using varying growth rates might enhance robustness or lead to different pattern types, such as interior stripe growth instead of the outer stripe addition shown in Fig 5D-middle [32]. Note that the growth rate used in this study is slower than the experimental growth rate in [12] as long times were simulated to reach convergence in the non-growing reflective boundaries case. Future research may want to test robustness for specific growth rates within an experimental system. Insights into which growth rates most effectively promote pattern formation could be valuable for optimizing experiments. Future work may also aim to develop a unified classification method that can interpret spatio-temporal patterns across different types of boundaries and both non-growing and growing domains. Such a method would produce comparable outputs to better understand the effects of growth and different boundary conditions on pattern formation. Although limited, our approach still identifies interesting cases where growth or boundaries influence patterning.

In conclusion, by including the realistic effects of multi-stability, growth, and boundary conditions, Turing patterns begin to bridge to biological applications. In particular, absorbing boundary conditions, by setting up chemical gradients reminiscent of Wolpert’s French flag model [33], improve robustness of the Turing parameter space significantly. As those effects generally require the full non-linear system and hence numerical treatment, this underscores the importance of looking beyond common linear stability analysis. Applications in engineering Turing patterns with synthetic circuits are a promising way forward.

## Methods

The 2-node non-linear Turing network given by Eq 1 is investigated using linear stability analysis and numerical methods. Numerically, the system is studied using non-growing domains with reflective boundary conditions, non-growing domains with absorbing boundary conditions, and finally growing domains with absorbing boundary conditions.

Numerical results are then classified into different types of patterns to understand the relationship between linear stability analysis and numerics, as well as the effects of boundary conditions and growth on patterning. The details of the methods mentioned are provided in this section. The following GitHub repository is available with the documented code necessary to produce the results shown in this paper: https://github.com/Endres-group/TuringNumerics_PLOS_2024

### Linear stability analysis

Linear stability analysis (LSA) is carried out to find out if a steady state exhibits a Turing instability (which is also called diffusion-driven instability). When it does, the system is capable of forming spatial patterns. As the name describes, diffusion-driven instabilities arise in these systems when a homogeneous steady state is stable to small perturbations in the absence of diffusion and becomes unstable in the presence of diffusion [34, 35].

The method of LSA, designed to check the system’s stability, will be explained for a generic two-morphogen reaction-diffusion system shown below:

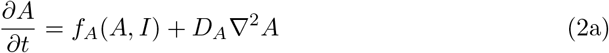

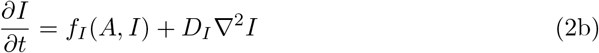

where *f*_*A*,*I*_ are the non-linear production terms and *D*_*A*,*I*_ are the diffusion constants of the two morphogens.

First, the steady states are defined *A*^∗^ and *I*^∗^, which satisfy the condition:

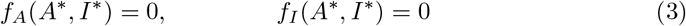

The non-linear reaction terms *f*_*X*_ (*A, I*) for *X* = *A, I* are then linearised around this steady state to investigate the instability to small perturbations around this steady state. Subsequently, the diffusion term *D*_*X*_ ∇^2^*X* is expressed as a cosine Fourier series which represents a solution with reflective boundary conditions. This results in the following expression:

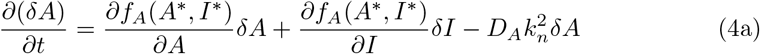

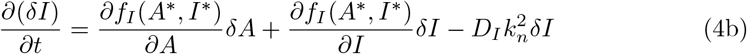

The reflective boundary conditions are ensured when the derivative of diffusion is zero at the boundaries *x* = [0, *L*]. Note that Dirichlet boundaries are not defined here as they are not studied using linear stability analysis, only numerically. With the constraints imposed by the reflective boundaries, *k*_*n*_ must be

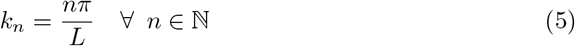

In this case, we are interested in the growth or decay of the perturbations over time. The stability of this system can be tested by calculating the eigenvalues *σ* of its Jacobian where matrix elements are evaluated at steady state

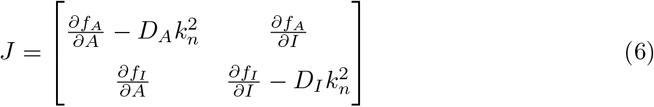

If *σ >* 0: perturbation (*δX*) grows making 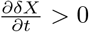. Therefore, the steady state is unstable. On the other hand, if *σ <* 0: perturbation (*δX*) decays making 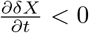. Therefore, the steady state is stable. *σ* = 0 is the marginal stable case, which generally requires further investigation, but numerically is easily avoided.

### Implementation of linear stability analysis

The steady states of the system are found using the Newton-Raphson algorithm with 100 initial conditions obtained using Latin-hypercube sampling with a uniform distribution ranging from [10^−3^, 10^3^]. The tolerance value of the Newton-Raphson algorithm is 10^−6^. The stability of these steady states is analyzed without diffusion by setting *k*_*n*_ = 0. If any of the eigenvalues have a real positive part, the steady state is unstable without diffusion. The stability of the steady state is analyzed by solving for the eigenvalues of the Jacobian (Eq 6) for all 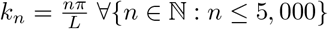,meaning 5,000 *k*’s are sampled to estimate the dispersion relation.

### Numerical methods

A finite difference method is chosen to solve the system of non-linear PDEs. By discretizing time and space, the two independent variables can be expressed as:

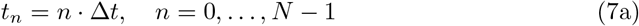

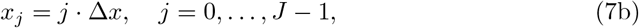

with *N* and *J* the number of discrete time and space points in our grid, respectively. Furthermore, Δ*t* and Δ*x* are the time steps and the space steps correspondingly, with *T* and *L* their final time and space values. The aim is to derive a numerical solution that approximates the unknown analytical solution, *U* (*j*Δ*x, n*Δ*t*) ≈ *u*(*j*Δ*x, n*Δ*t*), where *U* is the analytical solution and *u* is the numerical solution.

The Crank–Nicolson numerical scheme is chosen as it is unconditionally stable as shown by von Neumann stability analysis [36]. The unconditional stability is important to allow for larger Δ*t* and Δ*x*, without getting an amplification of errors. Larger Δ*t* and Δ*x* will result in reduced computational cost.

### Crank–Nicolson method

Consider a reaction-diffusion system with one spatial dimension

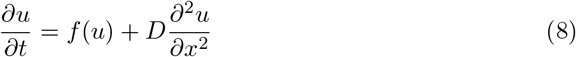

The spatial part of the equation can be approximated by

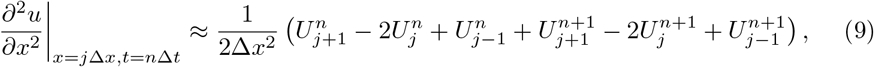

while the production function can be approximated to 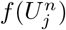. The approximations can be better visualized using the CN stencil (see Fig S1) Applying the CN stencil to the grid point (*i, j*), the reaction-diffusion system can be expressed as

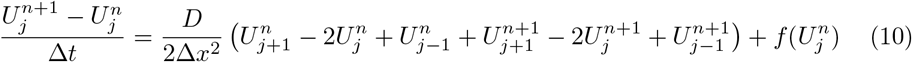

By reordering this approximation into a set of linear equations, the resulting problem is defined by matrices **A** and **B**, where 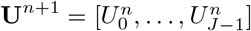 . The simplified system can be expressed as:

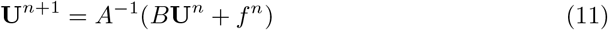

This method simplifies the complex system into a linear system that can be solved numerically. The returned solution is a 1D solution of the corresponding reaction-diffusion system. Although the method is unconditionally stable, the solution can contain oscillations if 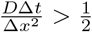[37]. Therefore, the ratio will be kept below 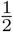 to avoid errors.

### Defining boundary conditions

The CN method can be implemented with Neumann reflective boundary conditions or Dirichlet absorbing boundary conditions. For Neumann reflective boundary conditions:

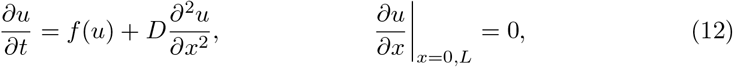

where the values of *U*_*j*_ at *j* = 0 and *j* = *J* − 1 are

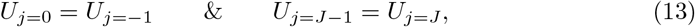

which are placed into the CN stencil shown in Eq 10. These values are chosen to ensure the derivative at the boundary is zero.

Similarly, Dirichlet absorbing boundary conditions are represented by the following system

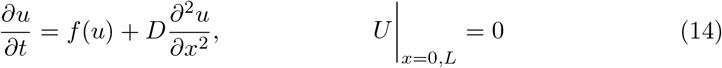

and have values of *U*_*j*_ at *j* = −1 and *j* = *J* such as

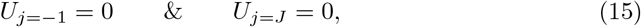

which are placed into the CN stencil shown in Eq 10, ensuring the value at the boundary is zero.

### Defining growing domains

Growth is introduced as apical isotropic linear growth, where cells are added to both boundaries with a linear growth rate. Linear growth is chosen as it is a good approximation to the growth observed in experimental bacterial colonies in Fig 1A [12]. In these colonies, cell division occurs mainly at the edges. Hence, we assume only edge division to simplify the model. To further reduce the model and computational power required, solutions are studied in 1D. Growth of the tissue is encoded in a 1D binary vector, where cells are denoted as 1 and empty spaces as 0. The number of 1’s grows linearly, which represents the expanding tissue. This vector is used as a mask, where 1’s determine the computation of reaction-diffusion terms and 0’s determine only the computation of diffusion.

### Classification methods

#### Reflective boundary classification

We first develop a method to classify the patterns produced numerically into (1) homogeneous, (2) temporal oscillator, as well as (3) non-stationary and (4) stationary patterns. We use a decision tree for the classification where the two layers are spatial homogeneity and convergence in time. This decision tree leads to the 4 types of patterns mentioned.

A pattern will be considered spatially homogeneous if the final snapshot *U* for any of the two molecular species fulfils the following condition

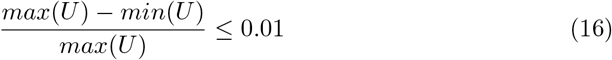

A pattern will be considered converged if the last 30 time points for any of the two molecular species fulfils the following condition

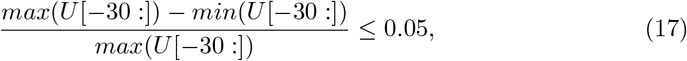

using common pseudocode splice notation for numerical arrays. The thresholds chosen were fine-tuned by testing them on the numerical patterns to obtain the best classification results. Using these two characteristics, spatial homogeneity and convergence, we can obtain 4 classes of patterns as seen in Fig S2:

1. Homogeneous patterns are homogeneous in space and converge in time.
2. Temporal oscillators, also called limit cycles, are homogeneous in space but do not converge, as they oscillate in time.
3. Non-stationary patterns are not homogeneous in space and do not converge in time.
4. Finally, stationary patterns are not homogeneous in space but converge in time.

#### Absorbing boundary classification

With absorbing boundary conditions and growth, patterns are rarely spatially homogeneous or converged, which makes the previous classification method unsuitable. Hence, a new classification system is developed based on the number of peaks. Peaks are detected using the Python *find peaks* algorithm with parameter prominence = 0.05.

Again, the thresholds for the peak finding algorithm need to be fine-tuned to avoid misclassification.

The peak classification shown in Fig S3, retrieves information on whether there is a pattern at all and whether this pattern is only a pattern at the boundary or a periodic pattern that scales with tissue length with constant wavelength as Turing patterns do. The 5 different types of patterns are:

1. Patterns with one peak are considered homogeneous as they result from the morphogens being reduced at the boundary due to absorption (Fig S3, no pattern, homogeneous).
2. Patterns with two peaks are also considered not to be patterned states as the two peaks might arise at the boundary for one of the diffusors due to the depletion of the other (Fig S3, no pattern, boundary effect).
3. Patterns with three peaks start displaying a pattern more similar to Turing repeats, although we cannot prove the number of peaks would scale with tissue length (Fig S3, weak pattern).
4. Patterns with four peaks could still be purely a boundary effect but this is less likely as the number of repeats points towards tissue scaling being possible (Fig S3, intermediate pattern).
5. Finally, patterns with five peaks and above are considered strong patterns and would be most similar to classical Turing patterns in non-growing reflective boundary domains (Fig S3, strong pattern).

## Supporting information

**Fig S1 Crank–Nicolson (CN) algorithm for numerical solution**. Geometric representation (stencil) with nodes and edges that represent the points of interest for the numerical approximation. The points of interest, which are represented by the equations in the Numerical Methods of the Methods section, are shown in green. Labels *j* and *n* are the current space and time points. The CN stencil has one spatial dimension and one temporal dimension, with axes labels time (*t*) and space (*x*).

**Fig S2 Decision tree for pattern classification in non-growing domains with reflective boundaries**. The decision tree is based on two layers: spatial homogeneity and convergence. The numerical solutions for the four different pattern outcomes include homogeneous, temporal oscillator, non-stationary, and stationary patterns, shown at the bottom. In the four numerical solutions, time is shown on the *y* axis, space on the *x* axis, and concentration by the color scheme.

**Fig S3 Pattern classification with absorbing boundaries**. Patterns are classified according to the number of peaks as shown in the 5 cases.

**Fig S4 Other types of multistable dynamics. (A)** Unstable state converges into Turing. **(B)** Unstable state produces pattern, while Turing state loses pattern. **(C)** Multistability disrupts all patterns. **(D)** Turing I Hopf state attracts the unstable state and generates a pattern.

**Fig S5 Confusion matrix for pattern transitions. (A)** Numerical outcome of reflective boundaries (*y* axis) versus absorbing boundaries (*x* axis).**(B)** Numerical outcome of non-growing domains (*y* axis) versus growing-domains (*x* axis). Numbers show the percentage of solutions across the row.

**Table S1 Kinetic parameters for PDE system**. A combination of the three values is used throughout this study to obtain a wide range of dynamical behaviors.

**Table S2 System parameters for numerical simulation**. Value 1 parameters are used for Figs 1, 2, 3, and 4, as well as Fig S2, Fig S3, and Fig S4. Value 2 parameters are used for Fig 5 and Fig S5.

## Acknowledgments

We gratefully acknowledge the support and funding provided by the EPSRC CDT BioDesign Engineering. We would also like to extend our sincere thanks to Prof. Mark Isalan for his invaluable guidance on the experimental aspects as well as Dr. Jure Tica and Dr. Dario Bazzoli for providing experimental images in Fig 1A,B. Additionally, we thank our colleagues, Dr. Roozbeh H. Pazuki and Antonio Matas Gil, for their insightful discussions.

