## Supplementary Material for "Effects of multistability, absorbing boundaries and growth on Turing pattern formation"

This supplementary material contains figures and tables with parameters which are referenced throughout the main text and supports the research findings.

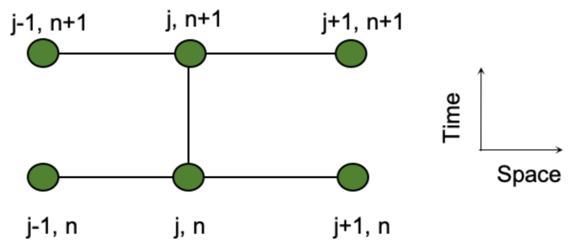


**Fig S1. Crank–Nicolson (CN) algorithm for numerical solution.** Geometric representation (stencil) with nodes and edges that represent the points of interest for the numerical approximation. The points of interest, which are represented by the equations in the Numerical Methods of the Methods section, are shown in green. Labels $j$ and $n$ are the current space and time points. The CN stencil has one spatial dimension and one temporal dimension, with axes labels time ($t$) and space ($x$).

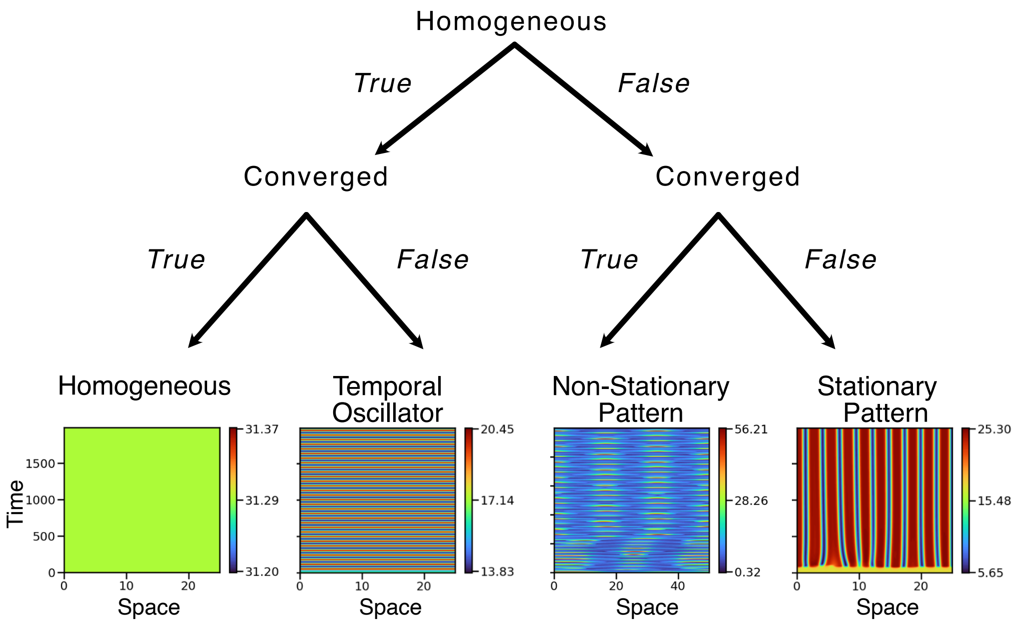


**Fig S2. Decision tree for pattern classification in non-growing domains with reflective boundaries**. The decision tree is based on two layers: spatial homogeneity and convergence. The numerical solutions for the four different pattern outcomes include homogeneous, temporal oscillator, non-stationary, and stationary patterns, shown at the bottom. In the four numerical solutions, time shown on the $y$ axis, space on the $x$ axis, and concentration by the color scheme.

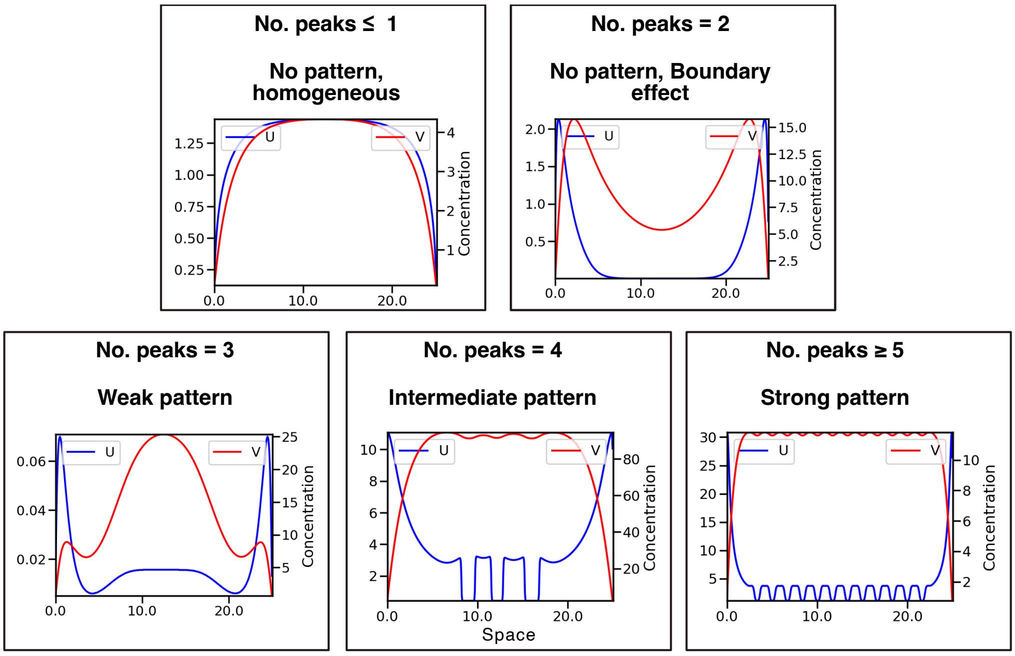


**Fig S3. Pattern classification with absorbing boundaries.** Patterns are classified according to the number of peaks as shown in the 5 cases.

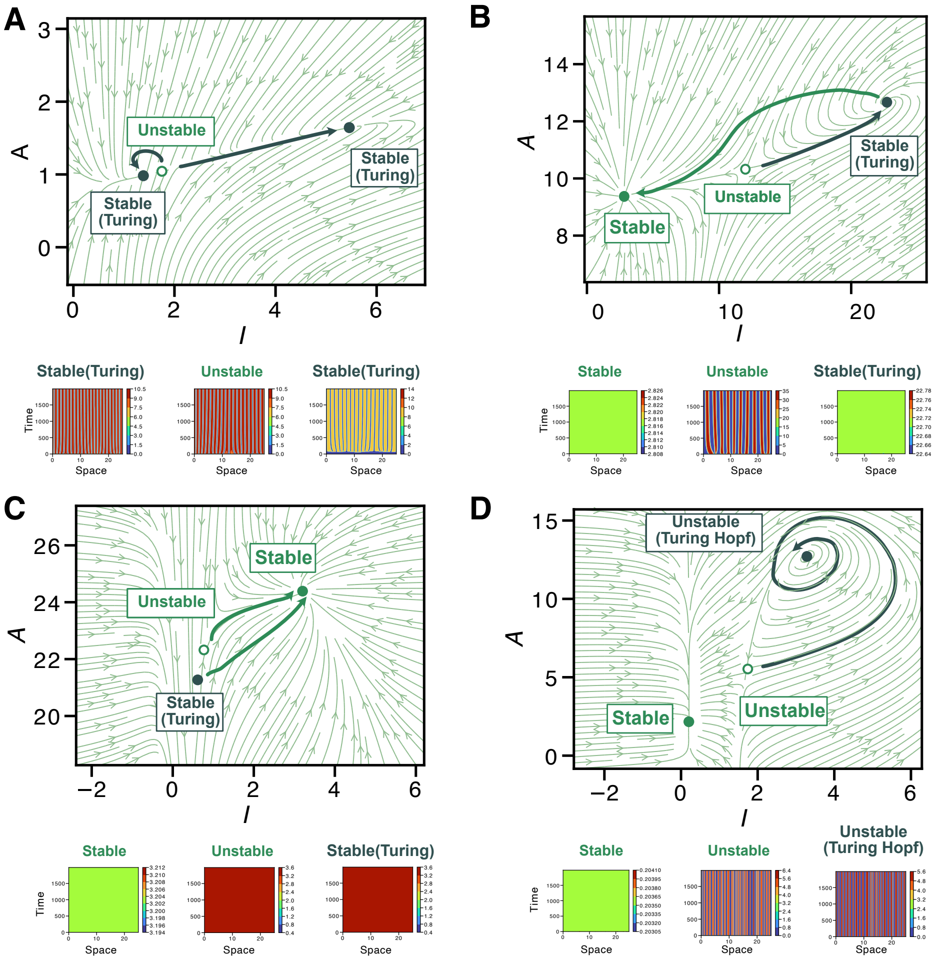


**Fig S4. Other types of multistable dynamics.** **(A)** Unstable state converges into Turing. **(B)** Unstable state produces pattern, while Turing state loses pattern. **(C)** Multistability disrupts all patterns. **(D)** Turing I Hopf state attracts the unstable state and generates a pattern.

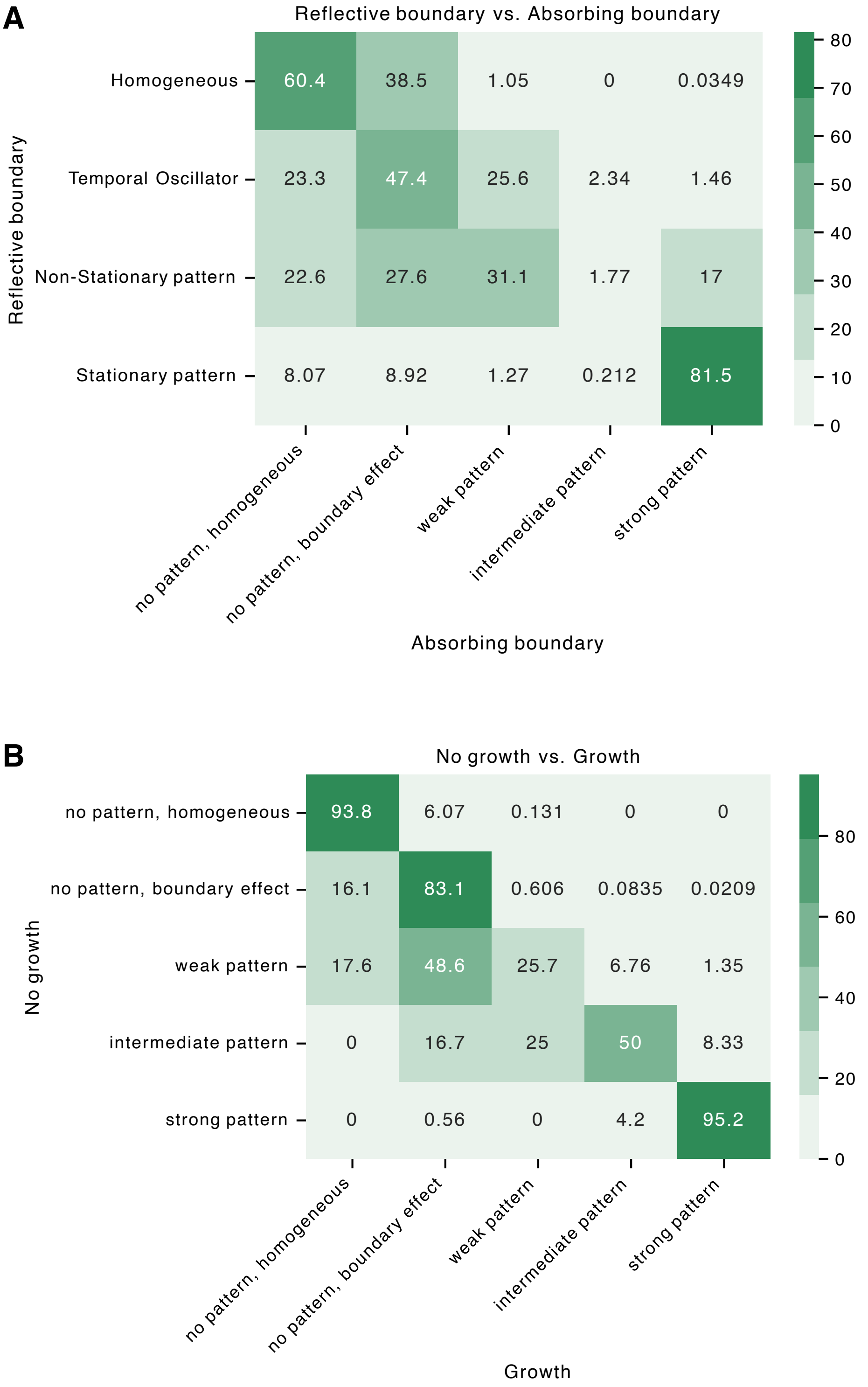


**Fig S5. Confusion matrix for pattern transitions. (A)** Numerical outcome of reflective boundaries ($y$ axis) versus absorbing boundaries ($x$ axis).**(B)** Numerical outcome of non-growing domains ($y$ axis) versus growing-domains ($x$ axis). Numbers show the percentage of solutions across the row.

| Parameter | Distribution | Value 1 | Value 2 | Value 3 |
|---|---|---|---|---|
| $V_m$ | Loguniform | (10, 1000) | (10, 10000) | (10, 10000) |
| $K_m$ | Loguniform | (0.1, 250) | (0.1, 100) | (0.1, 100) |
| $\mu_m$ | Loguniform | (0.001, 50) | (1, 100) | (1, 100) |
| $D_B$ | Loguniform / Fixed | (0.001, 10) | 0.01 | 10 |
| $D_A$ | Fixed | 1 | 10 | 0.01 |
| $b$ | Fixed / Loguniform | 0.01 | (10, 10000) | (10, 10000) |
| $n$ | Fixed / Loguniform | 2 | (2, 4) | (2, 4) |

**Table S1. Kinetic parameters for PDE system.** A combination of the three values is used throughout this study to obtain a wide range of dynamical behaviors.

| Parameter | $L$ | $\Delta x$ | $T$ | $\Delta t$ |
|---|---|---|---|---|
| **Value 1** | 25 | 0.05 | 2000 | 0.005 |
| **Value 2** | 100 | 0.2 | 18000 | 0.05 |

**Table S2. System parameters for numerical simulation.**Value 1 parameters are used for Figs. 1-4 as well as Figs. S2-S4. Value 2 parameters are used for Fig. 5 and Fig. S5.
